# Role of synaptic inhibition in the coupling of the respiratory rhythms that underlie eupnea and sigh behaviors

**DOI:** 10.1101/721829

**Authors:** Daniel S. Borrus, Gregory D. Conradi Smith, Christopher A. Del Negro

**Affiliations:** Department of Applied Science, Integrated Science Center, 540 Landrum Dr., William & Mary, Williamsburg, VA 23185

## Abstract

The preBötzinger Complex (preBötC) gives rise to two types of breathing behavior: eupnea and sighing. Here, we examine the neural mechanisms that couple their underlying rhythms by recording from the preBötC in neonatal mouse brainstem slice preparations. It has been proposed that chloride-mediated synaptic inhibition couples inspiratory (eupnea-related) bursts and sigh bursts, but we find no evidence to support that notion. First, we characterize a fluctuating temporal relationship between sigh bursts and their preceding inspiratory bursts; their coupling is far weaker than previously described. Surprisingly, selective blockade of inhibitory synapses strengthened (rather than weakened) that phasic inspiratory-sigh burst relationship. Furthermore, pharmacological disinhibition did not alter the duration of the prolonged interval that follows a sigh burst prior to resumption of the inspiratory rhythm. These results demonstrate that coupling between inspiratory and sigh rhythms does not depend on synaptic inhibition.

**SIGNIFICANCE STATEMENT:** Breathing consists of eupnea and sigh breaths, which differ in their magnitude and frequency. Both breath types emerge from a brainstem microcircuit that coordinates their timing. Here, we advance understanding of these rhythms by assessing the nature and strength of their coordination, and by showing that synaptic inhibition does not enforce their temporal coupling in contrast to conventional understanding. This study provides insights into the basic neural mechanisms that link oscillations of different amplitude and frequency in a core oscillator.

## INTRODUCTION

Breathing behavior consists of two interleaving rhythms and motor patterns: eupnea and sighing. Eupnea is the normal unlabored breathing that underlies periodic lung ventilation and drives alveolar gas exchange. Eupnea occurs at approximately 1-4 Hz in rodents (0.2-0.3 Hz in humans); each breath ventilates a small fraction of lung capacity. Sighs are also inspiratory breaths, but the volume of inhaled air during a sigh is two to five times that of a normal breath, and sigh frequency is an order of magnitude lower than the eupnea rhythm (Li and Yackle, 2017). Sighs reinflate collapsed (or collapsing) alveoli and are essential for optimal pulmonary function. Typically, sighs appear to ride atop ongoing eupneic breaths (Cherniack et al., 1981; Orem and Trotter, 1993), which suggests that periodically, but at a much lower frequency, the eupnea cycle triggers the sigh. After a sigh, the next eupneic breath is delayed for a duration roughly equivalent to one additional eupneic cycle (Cherniack et al., 1981; Orem and Trotter, 1993). This delay, which we refer to as the post-sigh apnea, suggests that sighs inhibit eupnea at least transiently. Eupnea and sigh breathing rhythms thus appear to be coupled, most likely via neural microcircuits of the brainstem that generate and control breathing movements.

In mammals, eupnea and sigh rhythms emanate from the preBötzinger Complex (preBötC) of the lower brainstem (Del Negro et al., 2018; Smith et al., 1991). Both rhythms are maintained in reduced slice preparations that isolate the preBötC as well as inspiratory premotor and motor neurons, and thus encapsulate a minimal breathing-related model system (Lieske et al., 2000; Ruangkittisakul et al., 2008; Chapuis et al., 2014). Because eupnea refers to behavior in living animals, *inspiratory* is the appropriate nomenclature for eupnea-related activity in slice preparations. Inspiratory rhythm depends on network properties, in which recurrent excitation among glutamatergic interneurons is rhythmogenic (Funk et al., 1993; Rekling et al., 2000; Del Negro et al., 2002; Wallen-Mackenzie et al., 2006; Feldman and Kam, 2015). The rhythmogenic mechanism of sighs is unknown, but it depends on neuropeptides released by parafacial respiratory interneurons (Li et al., 2016) as well as excitatory ionotropic and metabotropic receptor-mediated synaptic transmission (Lieske and Ramirez, 2006a, 2006b).

Inspiratory bursts appear to trigger sigh-related bursts, and in turn, sigh-related bursts delay the next inspiratory burst by almost an entire cycle (Lieske et al., 2000; Tryba et al., 2008). These observations *in vitro* mirror the *in vivo* coupling behavior described above, which suggests the mechanisms that couple inspiratory and sigh rhythms are contained within the preBötC and can be examined at the cellular and synaptic level *in vitro*.

What mechanisms couple these two rhythms? The only existing data suggest that glycinergic synaptic inhibition links the sigh-related burst to its preceding inspiratory burst, thus giving rise to the biphasic shape in which the inspiratory burst appears to trigger the sigh (Lieske et al., 2000). A recent mathematical model (Toporikova et al., 2015) posits two discrete systems for generating eupnea and sigh oscillations. The model inspiratory system acts on the sigh system via synaptic inhibition such that sigh bursts emerge via an escape-like process triggered by disinhibition at the tail end of inspiratory bursts. The model also suggests that the sigh system projects to the inspiratory system via fast excitatory synapses, and the strength of its excitation leads to a transient state of refractoriness, i.e., the post-sigh apnea, in the coupled system. However, the post-sigh apnea might also be attributable to synaptic inhibition from the sigh system onto the inspiratory rhythm generator.

Here, we test the role of synaptic inhibition in coupling eupnea and sigh rhythms *in vitro*. First, we elucidate the chloride reversal potential (E_Cl_) in order to verify that glycine and GABA_A_ synapses are inhibitory, and not functionally excitatory, as they are during embryonic development (Delpy et al., 2008; Ren and Greer, 2006). Next, we block glycinergic transmission and show that disinhibiting the preBötC *in vitro* does not uncouple the eupnea- and sigh-related rhythms, but in fact appears to couple them more strongly, given our observation of decreased latency between a sigh and its preceding inspiratory burst. We obtain similar results when we simultaneously block GABAergic and glycinergic transmission. We also show the duration of the post-sigh apnea does not depend on glycinergic or GABAergic transmission, which instead suggests that the post-sigh apnea reflects a refractory state attributable to post-synaptic membrane properties evoked by the sigh burst. These findings indicate that the eupnea and sigh rhythms are coupled predominantly by excitatory (rather than inhibitory) synaptic interactions.

## MATERIALS AND METHODS

### Ethical approval and animal use

The Institutional Animal Care and Use Committee at our institution approved these protocols, which conform to the policies of the Office of Laboratory Animal Welfare (National Institutes of Health, Bethesda, MD, USA) and the guidelines of the National Research Council (National Research Council (U.S.) et al., 2011). CD-1 mice (Charles River, Wilmington, MA) were maintained on a 14-hour light/10-hour dark cycle at 23° C and were fed *ad libitum* with free access to water.

Mouse pups of both sexes were anesthetized by hypothermia and then killed by thoracic transection at postnatal day 0 to 4. The neuraxis was removed in less than two minutes and further dissected in artificial cerebrospinal fluid (aCSF) containing (in mM): 124 NaCl, 3 KCl, 1.5 CaCl_2_, 1 MgSO_4_, 25 NaHCO_3_, 0.5 NaH_2_PO_4_, and 30 dextrose equilibrated with 95% O_2_-5% CO_2_, pH 7.4.

### Breathing-related measurements in vitro

Isolated neuraxes were glued to an agar block and then cut in the transverse plane to obtain a single 550-µm-thick slice that exposed the preBötC at its rostral face (Ruangkittisakul et al., 2011, 2014). Slices were then perfused with aCSF at 28° C in a recording chamber mounted below a stereomicroscope that enabled us to position suction electrodes under visual control. Extracellular K^+^ concentration ([K^+^]_o_) was increased to 9 mM to elevate preBötC excitability (Funk and Greer, 2013).

Inspiratory-related motor output was recorded from the hypoglossal (XII) nerve rootlets, which are captured in transverse slices along with the XII motoneurons and their axon projections to the nerve rootlets, using suction electrodes and a differential amplifier. Also, we simultaneously recorded field potentials from the preBötC by forming a seal over it with a suction electrode at the slice surface. Amplifier gain was set at 2000 and the band-pass filter was set at 300-1000 Hz. XII and preBötC bursts were full-wave rectified and smoothed for display and quantitative analyses of burst events. We acquired and digitized the signals at 4 kHz with a low-pass filter set to 1 kHz using a 16-bit analog-to-digital converter (ADInstruments, Colorado Springs, CO).

The glycine receptor antagonist strychnine hydrochloride (CAS number 1421-86-9, product S8753, Millipore Sigma, St. Louis, MO) and the GABA_A_ receptor antagonist picrotoxin (CAS number 124-87-8, product P1675, Millipore Sigma) were bath-applied at 5 µM while monitoring field potentials in the preBötC and XII motor output.

### Gramicidin patch recordings and glycine and muscimol application

We obtained whole-cell patch-clamp recordings under visual control from slices perfused in a recording chamber on a fixed-stage microscope (Zeiss AxioExaminer, Thornwood, NY). All recordings employed a HEKA EPC 10 patch-clamp amplifier (Holliston, MA). Patch pipettes were fabricated from borosilicate glass (OD: 1.5 mm, ID: 0.86 mm, 4-6 MΩ in bath) and filled with solution containing 150 mM KCl and 10 mM HEPES). We added gramicidin (CAS number 1405-97-6, product G0550000 from Millipore Sigma) acutely at the start of the experiment from stock solution (2 mg gramicidin per 1 ml dimethyl sulfoxide) such that the final concentration was 20 μg/ml. Patch pipettes were back-filled first with gramicidin-free patch solution in order to ensure a proper seal to the membrane. All recordings were corrected offline for a liquid junction of 3.74 mV (Barry and Lynch, 1991; Neher, 1992).

The experimental protocol began no earlier than 30 minutes after achieving a seal on the plasma membrane exceeding 1 GΩ (i.e., gigaohm seal), which was sufficient for gramicidin to form ionophores and thus allow intracellular access and current-clamp recording. We also fabricated pipettes (as described above) from which to eject glycine and muscimol (dubbed ‘puffer’ pipettes). Puffer pipettes were filled with 150 μM glycine and 30 μM muscimol diluted into ACSF (containing 9 mM [K^+^]_o_, as described above). 30 minutes after forming a gigaohm seal on the plasma membrane, the puffer pipette was positioned to within ∼50 µm from the neuron being recorded. 30 µM muscimol (CAS number 2763-96-4, product M1523, Millipore Sigma) and 150 µM glycine (CAS number 56-40-6, product 50046, Millipore Sigma) were ejected using 7-9 psi pressure pulses lasting 25-200 ms, which we triggered by TTL commands from the EPC-10 amplifier.

Cells were identified as neurons by their ability to discharge action potentials, recognizable ∼30 min after forming a gigaohm seal. Subsequently, we added 1 µM tetrodotoxin (TTX) to the bath to prevent spike-mediated chemical synaptic transmission. Neurons were held at a desired membrane potential using bias current. We measured transient changes in membrane potential in response to puffed glycine and muscimol, in which the previous 2 seconds of membrane voltage were used as baseline.

### Identification and categorization of sigh bursts

We distinguished a sigh burst from an inspiratory burst in field recordings based on burst magnitude, period regularity, and the presence of a post-sigh apnea. First, the area of sigh bursts in a given slice exceed the area of inspiratory bursts by more than one standard deviation away from the average area of all inspiratory bursts recorded in that slice preparation (because the frequency of sigh bursts is much lower than the frequency of inspiratory bursts, the average area of all bursts effectively returns the mean inspiratory burst area). Second, the cycle period for sigh bursts measures 1 to 4 minutes, and not outside this range (Lieske et al., 2000; Ruangkittisakul et al., 2008). Third, sigh bursts are followed by a prolonged inter-event interval (post-sigh apnea) greater than 1.3X the average inspiratory cycle time for six consecutive cycles preceding a putative sigh burst (Fig. 1). Burst events meeting two of these three conditions were considered sigh bursts (note that >90% of sigh bursts satisfied all three conditions).

**Figure 1.**
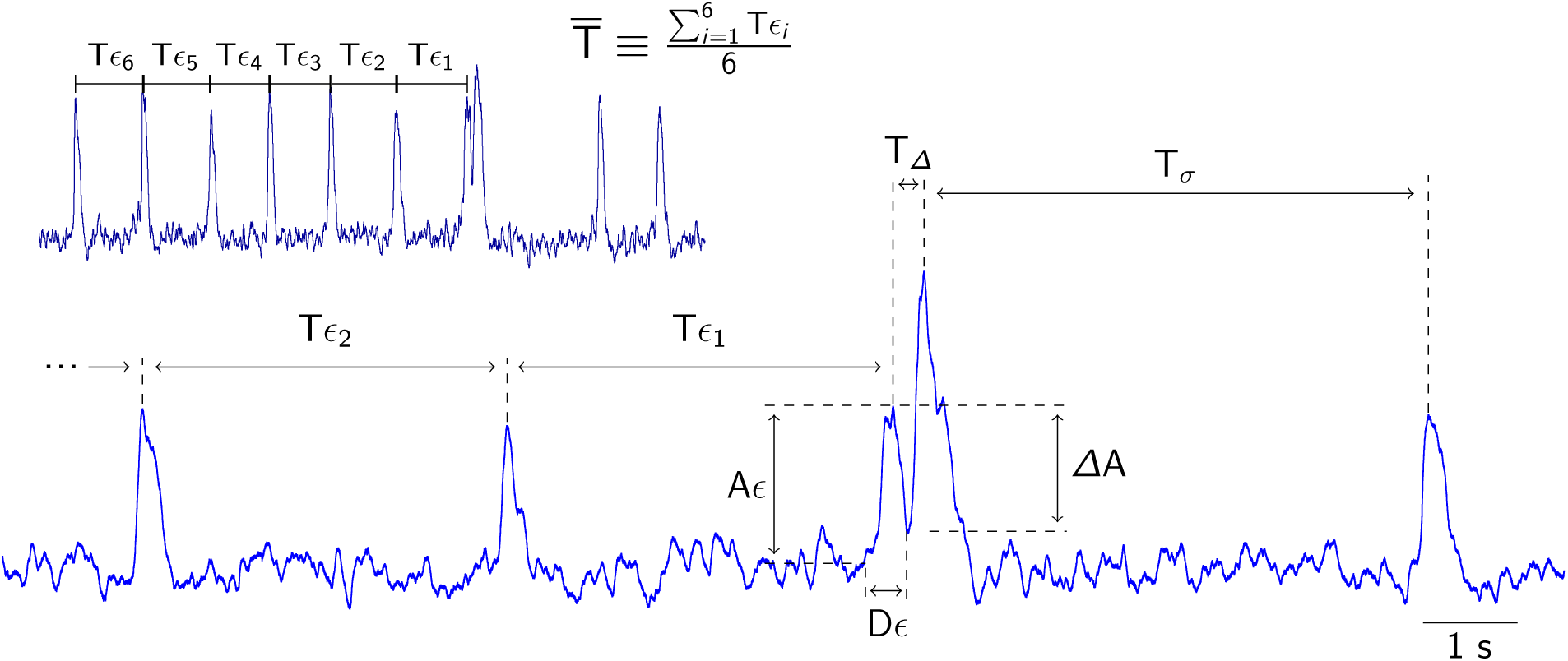
Integrated field recording of preBötC activity, which includes inspiratory and sigh bursts, in slices. Inset (upper left) shows normalized inspiratory cycle 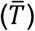 calculated from the average cycle period of the previous six inspiratory bursts 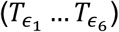. *T*_*ϵ*_ represents period of a single inspiratory cycle; *A*_*ϵ*_ represents amplitude of sigh-associated inspiratory burst; Δ*A* reflects voltage drop from peak of an inspiratory burst to the nadir during the inspiratory-sigh interval; *D*_*ϵ*_ reflects the duration of sigh-associated inspiratory burst; *T*_*Δ*_ represents the inspiratory-sigh interval; and *T*_σ_ reflects the duration of the post-sigh apnea.

We used an algorithmic rule for categorizing sigh bursts into the five characteristic types: *long interval, doublet, classic, S to I*, and *conjoint*. Figure 1 provides an accompanying schematic. Epsilon (*ϵ*) indicates eupnea-related inspiratory cycles. Sigma (*σ*) indicates sigh-related cycles. Delta (Δ) indicates time or amplitude differences. For each sigh burst, we computed six representative metrics. The average inspiratory cycle time 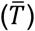 is the average inspiratory cycle time, computed from six inspiratory cycles preceding the sigh 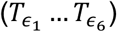. *T*_Δ_ is the inspiratory-sigh interval computed as the peak time of the sigh burst minus the peak time of the inspiratory burst. *T*_*σ*_ is the duration of the post-sigh apnea. For each sigh burst, the duration of the associated inspiratory burst is denoted *D*_*ϵ*_. The peak amplitude of that inspiratory burst is *A*_*ϵ*_, and the amplitude of the voltage drop from the peak of the preceding inspiratory burst to the trough of the inspiratory-sigh interval is Δ*A*. Long interval sigh bursts are defined by 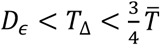, doublet sighs are defined by *D*_*ϵ*_ > *T*_Δ_ and 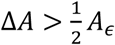, classic sighs are defined by 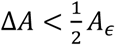, S-to-I sigh bursts are defined by *T*_Δ_ < 0 (because the inspiratory burst follows the sigh burst); and conjoint sighs are defined by 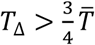

### Measurements and statistics

We measured the amplitude (V) and area (V-s) of eupnea-related inspiratory bursts (frequency range 0.1–0.2 Hz) and lower-frequency sighs (<1 min^-1^). Burst amplitude was measured from baseline (defined as the average field potential 500 ms prior to the burst) to the burst peak. In all cases inspiratory and sigh burst peaks are distinct and distinguishable. Nevertheless, for classic, S-to-I, conjoint bursts, and some doublet bursts, the amplitude of the trailing burst is enhanced by temporal summation after the preceding burst.

Burst area was computed as the area under the curve of the trajectory of the field potential recording. For classic, S-to-I, conjoint bursts, and some doublet bursts, the burst area measurement conflates the two partially coincident events. In such cases, the aggregate area is classified sigh burst area. For long interval sighs – to be consistent with the other categories described above – the area of the sigh burst was always summed with the area of the preceding inspiratory burst regardless of the inspiratory-sigh interval duration.

We analyzed data and computed statistics using LabChart 7 (ADInstruments), MATLAB 2018b (Mathworks, Natick MA), and Igor Pro 8 (Wavemetrics, Oswego, OR). We describe the statistical hypothesis tests used as they appear in the main text.

## RESULTS

### Chloride-mediated fast synaptic drive in the preBötC of neonatal mice is inhibitory

We recorded preBötC neurons through gramicidin-perforated patches selectively permeable to monovalent cations. The internal chloride concentration remains unchanged so E_Cl_ can be determined (Kyrozis and Reichling, 1995). In the presence of TTX, pressure pulse ejections of muscimol and glycine transiently perturbed the membrane potential, which reversed at or below –45 mV (Fig. 2A and B). E_Cl_ measured –49 ± 8 mV (mean ± SD, 7 preBötC neurons recorded in 6 slices). We observed no relationship between E_Cl_ and age (Fig. 2C), which indicates that E_Cl_ is consistent during early postnatal development.

**Figure 2.**
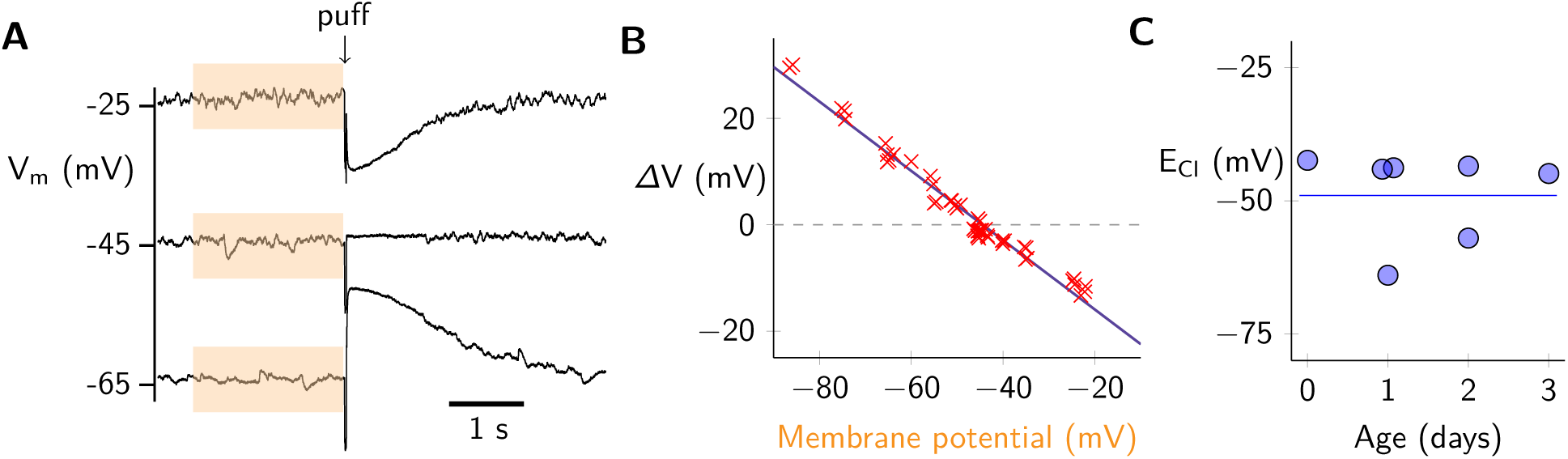
Determining the chloride reversal potential (E_Cl_) in preBötC neurons in neonatal brainstems. A: Representative traces of membrane voltage before and after puffer application of 150 μM glycine and 30 μM muscimol. Mean membrane potential before puffer application was calculated from the previous 2 seconds (orange box) of recording. Change in membrane voltage (Δ mV) was calculated as the voltage change from mean membrane potential to peak membrane response. B: Membrane potential changes in response to glycine and muscimol puffs plotted versus holding potential from a single representative experiment (same cell as A). E_Cl_ (Δ mV = 0) is calculated using a linear regression (purple line). C: E_Cl_ from 7 cells (N = 6 mice) plotted as a function of postnatal mouse age. Dashed line indicates the average E_Cl_ for the sample.

### Sigh bursts have a variable inspiratory-sigh interval

We measured 343 sigh bursts in 13 slice preparations (26 ± 7 sigh bursts per slice). The classic sigh pattern, which is often described as the sigh burst building off the crest of an inspiratory burst, does not accurately describe the temporal relationship between most sigh bursts and their preceding inspiratory bursts. To illustrate the variability of the inspiratory-sigh intervals we identified five discrete categories (Fig. 3A; the algorithm we used to assign sigh bursts to one of these five categories is explained in Materials and Methods). The classic biphasic sigh burst was only recorded 72 times (∼21%). Sigh bursts with long intervals and doublets were observed 97 times (∼28%) and 56 times (∼16%), respectively.

**Figure 3.**
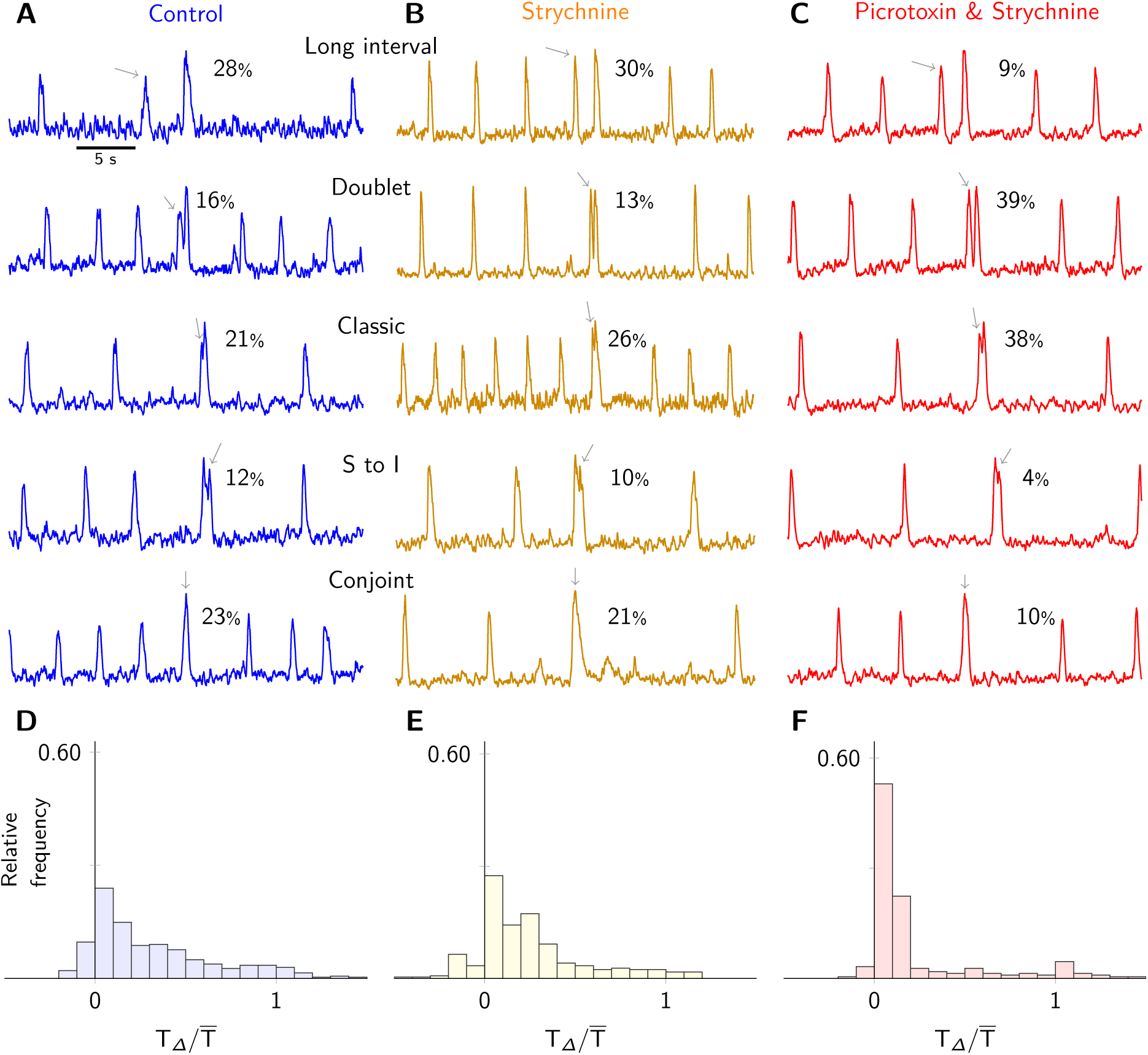
The variability in timing of sigh bursts. A-C: Integrated field recordings of preBötC activity showing the five variations of the inspiratory-to-sigh burst coupling during control (A), and after treatment with 5 μM strychnine (B), as well as treatment with 5 μM strychnine and 5 μM picrotoxin (C). Percentages next to sigh categories display the fraction a particular sigh class was observed out of all sighs recorded in that condition (343 control sighs, 292 strychnine sighs, 219 strychnine & picrotoxin sighs). Grey arrows identify the sigh-associated inspiratory burst. D-F: Histograms of the normalized inspiratory-sigh interval 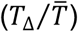 under all three conditions. The width of each bar is 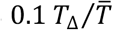.

We also observed 40 sigh bursts immediately followed by an inspiratory burst, which results in a negative inspiratory-sigh interval (∼12%), dubbed S-to-I sigh bursts (Fig. 3A). To our knowledge, the S-to-I phenomenon has not previously been experimentally documented, and it suggests that a sigh burst does not absolutely require a preceding inspiratory burst to trigger it.

Further, we recorded 78 conjoint sigh bursts (∼23%) in which the inspiratory and sigh bursts appear to occur simultaneously. This interpretation cannot be confirmed because it is possible that the sigh burst we label as conjoint actually occurs with a delay equivalent to an entire inspiratory cycle time. In the other cases shown in Fig. 3A, the delay from the inspiratory to the sigh burst is less than the average inspiratory cycle time so one can perceive their linkage. However, if the conjoint sigh bursts were actually isolated sighs – independent of and separated from preceding inspiratory bursts by an entire cycle – then their amplitude and area measurements should be demonstrably smaller than the amplitude and area of all other categories of sigh bursts. However, the mean burst amplitude of putative conjoint sigh bursts (34 ± 12 mV) was commensurate with the mean amplitude of all other sigh burst types (36 ± 13 mV) (Wilcoxon rank sum test, p = 0.73, n = 9 slices). Similarly, the mean burst area of putative conjoint sigh bursts (18 ± 10 mV-s) and all other sigh burst types (20 ± 11 mV-s) were also commensurate (Wilcoxon rank sum test, p = 0.26, n = 9 slices). This suggests that the events in question consist of synchronized inspiratory and sigh bursts (i.e., conjoint bursts).

Blocking glycinergic synapses with strychnine did not alter the relative prevalence of inspiratory-sigh interval categories (Fig. 3B). However, after blocking all chloride-mediated ionotropic synaptic receptors with strychnine and picrotoxin, it appeared that doublet and classic sighs increased by approximately two-fold or more (13% to 39% and 26% to 38%, respectively), even though we continued to observe all of the inspiratory-sigh interval categories (Fig. 3C). These data show that the temporal coupling between inspiratory and sigh bursts is more variable than previously reported. Furthermore, removal of chloride-mediated inhibition may favor sigh bursts in which temporal coupling is the tightest, i.e., the doublet and classic types of sigh bursts.

### Inhibitory synapses do not couple inspiratory and sigh bursts

In order to quantitatively compare inspiratory-sigh intervals, we calculated their probability distributions (Fig. 3D-F) and cumulative distribution functions (Fig. 4). In control conditions, a sigh burst was most likely to manifest at relatively short intervals following an inspiratory burst. This is illustrated by the peak probability of a sigh burst at the earliest part of the normalized inspiratory cycle; the probability then tapers off at later stages of the normalized cycle. Indeed, 24% of sigh bursts occur within the first tenth of the normalized cycle time and the majority (64%) occur within the first half of the normalized cycle (Fig. 3D).

**Figure 4.**
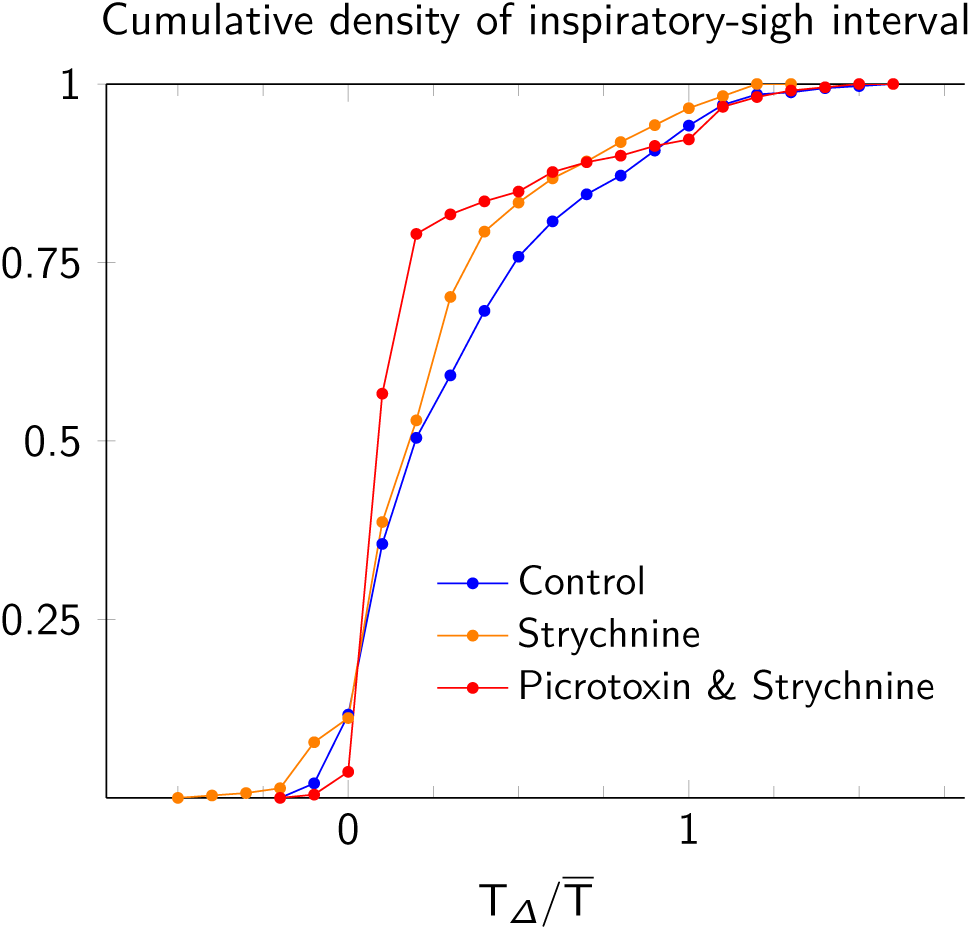
Cumulative density distribution of the normalized inspiratory-sigh interval 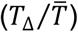) under control conditions (blue), treatment with 5 μM strychnine (orange), and treatment with both 5 μM strychnine and 5 μM picrotoxin (red).

Using strychnine to block glycinergic synaptic transmission, the probability distribution of inspiratory-sigh intervals remained weighted towards the first half of the normalized cycle (Fig. 3E), suggesting that inspiratory-sigh coupling remained intact. In contrast, previous studies suggested that blocking glycinergic synapses removed any temporal relationship between the sigh burst and its preceding inspiratory burst (Lieske et al., 2000; Chapuis et al., 2014), in which case the probability distribution in Fig. 3E would be uniformly distributed between 0 and 1. Here, the probability of short inspiratory-sigh intervals 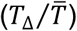 increased such that the average normalized inspiratory-sigh interval decreased from 0.31 ± 0.34 in control to 0.25 ± 0.30 in the presence of strychnine. This trend is reflected in the significant leftward shift of the cumulative distribution function from control to treatment with strychnine for cycle time > 0 (Fig. 4, Kolmogorov-Smirnov, test statistic = 0.13, p = 0.011, n = 7 slices).

Similarly, when we simultaneously blocked glycinergic and GABA_A_ receptors with a strychnine and picrotoxin cocktail, the sigh burst coupled more tightly with the preceding inspiratory burst than in control conditions. The probability of short inspiratory-sigh intervals 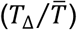 increased from control to the strychnine and picrotoxin cocktail such that the average normalized inspiratory-sigh interval decreased from 0.31 ± 0.34 in control to 0.21 ± 0.32 in strychnine and picrotoxin. The significant leftward shift of the cumulative distribution function for cycle time > 0 further demonstrates that when both glycinergic and GABA_A_ receptors were blocked, the sigh burst was more likely to occur earlier with respect to the preceding inspiratory burst than during control (Fig. 4, Kolmogorov-Smirnov, test statistic = 0.31, p = 4.0E-12, n = 6 slices). It is worth noting that the large standard deviation of the average normalized inspiratory-sigh interval, relative to the mean interval in all conditions, reflects the variability in the timing of a sigh discussed earlier and illustrated in Figure 3A-C.

Our analyses (Figs. 3 and 4) show the removal of chloride-mediated synaptic inhibition does not uncouple the sigh from its preceding inspiratory burst, rather it strengthened the temporal coupling of inspiratory and sigh bursts.

### Inhibitory synapses do not influence post-sigh apnea

We calculated the relative post-sigh apnea as the duration of the post-sigh interval (*T*_*σ*_) divided by the average inspiratory cycle time 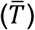, thus normalizing it and expressing it as a unitless ratio. Then, we compared the relative post-sigh apnea in control and after blocking either glycinergic transmission, or both glycinergic and ionotropic GABAergic transmission (Fig. 5).

**Figure 5.**
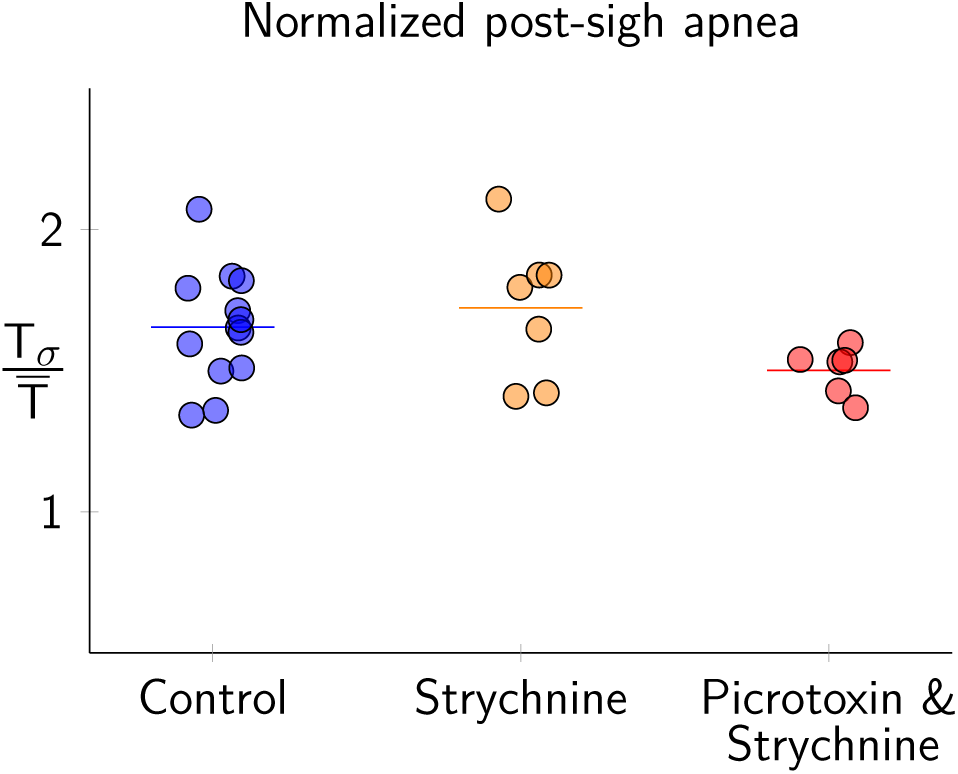
Average post-sigh apnea duration normalized by the average inspiratory cycle time 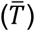 for control conditions (blue), treatment with 5 μM strychnine (orange), and treatment with both 5 μM strychnine and 5 μM picrotoxin (red). Each dot represents the average normalized post-sigh apnea for an experiment. Horizontal bars represent the average of all experiments.

The relative post-sigh apnea measured 1.65 ± 0.20 in control, 1.72 ± 0.25 in strychnine, and 1.5 ± 0.09 in the strychnine-picrotoxin cocktail. These measurements are statistically indistinguishable (one-way ANOVA, test statistic = 2.12, p = 0.14, n = 13 slices).

## DISCUSSION

Inspiratory and sigh-related rhythms both emerge from the preBötC. Their interactions and influence on one another can be studied *in vitro*. The conventional understanding is that sigh bursts build off the crest of inspiratory bursts, which implies strong coupling wherein inspiratory bursts trigger sigh bursts. However, that conceptual framework oversimplifies how sigh bursts actually emerge from preBötC activity. There is previously unrecognized variability in the timing between inspiratory and sigh bursts, which suggests that their coupling is weaker, relatively speaking, than previously appreciated. The existence of S-to-I and conjoint sigh bursts further reinforces this notion of flexibility in the relationship between a sigh burst and an associated inspiratory burst by showing that an inspiratory burst is not strictly necessary to trigger a sigh burst. Nevertheless, more often than not an inspiratory burst does trigger a sigh burst with an interval whose duration is on the order of one-tenth of the average inspiratory cycle time.

Here, in early postnatal mouse development (P0-4) with elevated (9 mM) [K^+^]_o_ ACSF to boost slice excitability, E_Cl_ measured –49 mV. This Nernst equilibrium potential is below spike threshold and approximates the level of baseline membrane potential during the interburst interval. Therefore, GABA_A_ and glycinergic inputs either shunt the membrane – rendering it less responsive to excitatory (depolarizing) drive – or hyperpolarize it directly during the preinspiratory phase or during the inspiratory burst itself when the membrane potential trajectory exceeds –49 mV.

We show that chloride-mediated synaptic inhibition is not responsible for the temporal coupling between the inspiratory and sigh bursts. Rather, disinhibition strengthened their coupling. Therefore, our primary conclusion is that excitatory (not inhibitory) synaptic transmission links the inspiratory and sigh rhythms of the preBötC.

This conclusion contradicts prior studies showing that blockade of glycinergic transmission decoupled sighs from their preceding inspiratory bursts and created free-running sigh burst rhythms that appeared to be independent from ongoing inspiratory rhythms (Lieske et al., 2000; Chapuis et al., 2014; Toporikova et al., 2015).

The discrepancy between those prior results and our present findings are probably attributable to the late embryonic reversal of the chloride electrochemical gradient. Before embryonic day 15.5 (E15.5) in mice, the dominant expression of cotransporter NKCC1 in brainstem and spinal cord neurons elevates intracellular chloride concentration (Delpy et al., 2008; Ren and Greer, 2006; Viemari et al., 2011) such that chloride-mediated synaptic currents are inward (i.e., excitatory) at the baseline membrane potential of rhythmically active preBötC interneurons. Perinatally NKCC1 expression decreases in parallel with increasing expression of the chloride symporter, KCC2, which lowers intracellular chloride concentration. In the mature state, dominant KCC2 expression ensures that the chloride equilibrium potential is more hyperpolarized than spike threshold as well as baseline membrane potential during rhythmic activity. The mature gradients ensure that chloride-mediated synaptic currents are outward and inhibitory.

Whereas we studied postnatal (P0-4) mice exclusively, Chapuis et al. (2014) and Toporikova et al. (2015) studied embryonic mice (E15.5-18.5). At these ages, with elevated (8 mM) [K^+^]_o_ ACSF, E_Cl_ is above spike threshold and consequently glycinergic synapses serve to depolarize and evoke action potentials in preBötC neurons (Ren and Greer, 2006; Delpy et al., 2008). Under these conditions, we infer that glutamatergic, glycinergic and GABA_A_ergic synapses are effectively excitatory and link sigh bursts to their preceding inspiratory bursts. When net excitatory drive is perturbed (such as by blocking chloride-mediated synaptic excitation) then inspiratory-sigh coupling weakens, and the sigh rhythm appears to ‘free run’ in a manner that is uncoupled from inspiratory rhythm.

Development and chloride gradients might also explain the discrepancies between our results and Lieske et al. (2000), who also concluded that glycinergic synapses couple inspiratory and sigh rhythms. Those authors reported using mice aged 0-2 weeks, a postnatal window that overlaps and extends beyond ours. We surmise that their sigh burst experiments were performed exclusively or predominantly using preparations from P0 mice with immature chloride gradients. Then the same explanation holds; chloride gradients favoring inward currents (with suprathreshold reversal potential) render glycine synapses ostensibly excitatory.

Chloride-mediated synaptic inhibition does not contribute to the post-sigh apnea. Instead, the post-sigh apnea is caused by activation of the intrinsic cellular mechanisms that help terminate inspiratory bursts, which are recruited to a greater degree during sigh events (compared to typical inspiratory cycles). These burst-terminating mechanisms include activity-dependent outward currents such as the electrogenic Na/K ATPase pump current, Na^+^-dependent K^+^ current, and ATP-dependent K^+^ current (Del Negro et al., 2009; Krey et al., 2010), as well as excitatory synaptic depression (Kottick and Del Negro, 2015; Guerrier et al., 2015). It was recently shown that the magnitude of inspiratory burst-related depolarization directly evokes corresponding levels of post-burst hyperpolarization in preBötC neurons, from which the neurons must recover prior to generating the next inspiratory burst (Baertsch et al., 2018). The sigh burst in this context is an extreme version of that same mechanism: the increased magnitude and duration of the sigh event correspondingly evokes larger-than-average activity-dependent refractory (outward) currents and depresses excitatory synapses to a greater extent than during typical inspiratory bursts of lower magnitude and duration. The larger-than-average hyperpolarization (and depressed synapses) extends the duration of the interburst interval, thus creating the post-sigh apnea.

Here we demonstrate that chloride-mediated synaptic communication is inhibitory in the preBötC of neonatal mice. Although sigh bursts are often closely preceded by inspiratory bursts, their temporal coordination is more variable than previously documented. Chloride-mediated synaptic inhibition plays no obligatory role in coupling the inspiratory and sigh rhythms in postnatal mice and we speculate that this principle may hold for juvenile and adult stages of development because E_Cl_ is expected to remain below spike threshold and inhibitory; in fact we expect it to descend lower than –49 mV during further maturation. A model of the preBötC core that incorporates inspiratory and sigh oscillators need not include synaptic inhibition to generate or couple the discrete rhythms. This emphasis on the primacy of excitatory synaptic interactions probably extends to embryonic development when chloride-mediated synapses are functionally excitatory.

